# Intermittent energy restriction inhibits tumor growth and enhances paclitaxel response in a transgenic mouse model of endometrial cancer

**DOI:** 10.1101/2024.02.02.578679

**Authors:** Ziyi Zhao, Jiandong Wang, Weimin Kong, Ziwei Fang, Michael Coleman, Ginger Milne, Wesley C. Burkett, Meredith A. Newton, Douglas Lee, Beor Deng, Xiaochang Shen, Hongyan Suo, Wenchuan Sun, Stephen Hursting, Chunxiao Zhou, Victoria L Bae-Jump

**Author notes:** Both authors contributed equally to this work (Ziyi Zhao, Jiandong Wang).

## Abstract

**Objective:** Overweight/obesity is the strongest risk factor for endometrial cancer (EC), and weight management can reduce that risk and improve survival. We aimed to establish the differential abilities of intermittent energy restriction (IER) and low-fat diet (LFD), alone and in combination with paclitaxel, to reverse the procancer effects of high-fat diet (HFD)-induced obesity in a mouse model of EC.

**Methods:** *Lkb1*^*fl/fl*^*p53*^*fl/fl*^ mice were fed high-fat diet (HFD) or LFD to generate obese and lean phenotypes, respectively. Obese mice were maintained on HFD or switched to LFD (HFD-LFD) or IER (HFD-IER). Ten weeks after induction of endometrial tumor, mice in each group received paclitaxel or placebo for 4 weeks. Body and tumor weights; tumoral transcriptomic, metabolomic and oxylipin profiles; and serum metabolic hormones and chemocytokines were assessed.

**Results:** HFD-IER and HFD-LFD, relative to HFD, reduced body weight; reversed obesity-induced alterations in serum insulin, leptin and inflammatory factors; and decreased tumor incidence and mass, often to levels emulating those associated with continuous LFD. Concurrent paclitaxel, versus placebo, enhanced tumor suppression in each group, with greatest benefit in HFD-IER. The diets produced distinct tumoral gene expression and metabolic profiles, with HFD-IER associated with a more favorable (antitumor) metabolic and inflammatory environment.

**Conclusion:** In *Lkb1*^*fl/fl*^*p53*^*fl/fl*^ mice, IER is generally more effective than LFD in promoting weight loss, inhibiting obesity-related endometrial tumor growth (particularly in combination with paclitaxel), and reversing detrimental obesity-related metabolic effects. These findings lay the foundation for further investigations of IER as a EC prevention strategy in women with overweight/obesity.

## Introduction

An estimated 65,950 women in the United States were projected to be diagnosed with endometrial cancer (EC) in 2023, and 12,550 to succumb to the disease, making it the most prevalent gynecological cancer(1). Among the risk factors for the development of EC, including but not limited to increasing age, physical inactivity, and diabetes mellitus, obesity is the strongest, with approximately 57% of cases in the United States linked to being overweight/obese (2). Unfortunately, The incidence and mortality of EC is steadily increasing in the United States due to the aging population and increasing prevalence of obesity (3). While most EC cases are diagnosed at an early stage and have a favorable prognosis, with survival rates of up to 80%, people with high-risk or advanced disease at diagnosis have a poor prognosis with few effective treatment strategies(4). Thus, there is a pressing need to improve our understanding of the obesity-related mechanisms underlying EC and to develop new preventive strategies and treatment regimens to improve prognosis in patients with EC.

Limited data suggests women who have experienced weight loss at some point during adulthood tend to have a lower risk of developing EC compared to those who have not lost weight (5, 6). Dietary interventions involving energy restriction, including chronic calorie restriction (CR) and intermittent energy restriction (IER), have demonstrated various physiological advantages, such as weight loss, decreased inflammation, enhanced insulin sensitivity, and modifications to the gut microbiome in humans(7). Consistently across animal studies, including those in rodents and monkeys, chronic CR without malnutrition is a potent cancer prevention strategy, effectively preventing spontaneous and chemically induced tumors such as lymphomas, mammary cancers, bladder cancer and skin cancers(7-9). Patients with obesity-related cancers, such as EC, may experience more anticancer advantages from CR than those with non-obesity-related cancers(10). Due to CR being challenging to maintain long-term and IER being more tolerable and sustainable, IER has become a popular alternative to CR for studying cancer prevention and treatment in animals and humans (5). CR and IER are equally effective in decreasing body weight and fat mass in individuals who are overweight or obese, but IER regimens may yield slightly more favorable retention of lean mass(11).

Several studies have demonstrated that IER prevents or delays tumor growth in multiple animal models(12, 13). However, the relative abilities of IER and low-fat diet (LFD; another dietary weight loss intervention) to promote weight loss, mitigate high-fat diet (HFD)-induced procancer perturbations, and reduce obesity-driven tumor burden remain unclear in models of EC. Furthermore, the antitumor benefit of combining paclitaxel (PTX), a common chemotherapy, with these dietary interventions is unknown. In this study, using a transgenic mouse model of EC, we aimed to establish the differential anti-obesity and anticancer effects of IER and LFD, with and without paclitaxel PTX, relative to HFD, and compare the tumoral transcriptional, metabolomic, and oxylipin profiles and systemic hormone and chemocytokine levels of obese, phenotypically formerly obese, and lean mice.

## Materials and methods

### Diet interventions in Lkb1^fl/fl^p53^fl/fl^ mouse model of EC

This study used a female *Lkb1*^*fl/fl*^*p53*^*fl/fl*^ mouse model developed in our laboratory and shown to develop endometrioid EC following induction (14). The animal protocol (# 21-229) for this study was approved by the Institutional Animal Care & Use Committee of the University of North Carolina at Chapel Hill (UNC-CH). All mice were housed in a 12-hour light/dark cycle with a temperature of 22±2°C. Mice were fed a high-fat diet (HFD, 60 kcal% fat, D12492; Research Diets, New Brunswick, NJ), or a low-fat diet (LFD, 10 kcal% Fat, D12450J; Research Diets) from 3 weeks of age to generate obese and lean phenotypes, respectively. At 12 weeks of age, the mice fed HFD were randomly assigned to one of three groups: continue HFD, switch to LFD (HFD-LFD), or switch to IER (HFD-IER). At 16 weeks of age, mice were injected with 5 µl adenovirus expressing Cre recombinase (AdCre; University of Iowa Transfer Vector Core, a titer of 10^11^-10^12^) into the left uterine horn to induce EC. At 26 weeks of age, mice were randomly divided into eight groups (30-35 mice/group): HFD; HFD+PTX; LFD; LFD+PTX; HFD-LFD; HFD-LFD+PTX; HFD-IER; HFD-IER+PTX. PTX or vehicle (injected in all mice that didn’t receive PTX) was administered for 4 weeks. At 30 weeks of age, all mice were euthanized. Blood was collected and processed to obtain serum, which was stored at -80°C prior to assay. Gonadal adipose pads and endometrial tumors were collected and weighed; half of each tumor as formalin-fixed for histopathology and half stored in liquid nitrogen until assayed. For the HFD-IER and HFD-IER+PTX groups, the amount of food given provided 70% of the average kilocalories consumed by the mice fed LFD during the previous week. The IER regimen involved a 13% CR diet (D15032803, Research Diet) for 5 days/ week and a 70% CR diet (D15032804, Research Diet) for 2 non-consecutive days/week(15, 16). Body weights were measured daily.

### Serum hormones, cytokines, and chemokines

Serum levels of hormones, inflammatory cytokines and chemokines were measured using the mouse multiplexed Luminex assays (Bio-Rad, Hercules, CA) according to the manufacturer’s protocols in the University of North Carolina at Chapel Hill (UNC-CH) Animal Core Facility. Data was collected and analyzed using the Luminex-200 system (Austin, TX).

### Tumor immunohistochemistry (IHC)

EC slides (4 µm) from the *Lkb1*^*fl/fl*^ *p53*^*fl/fl*^ mice were de-paraffinized, and recovery of antigen reactivity of the slides was achieved using heat-induced epitope retrieval. Ki67, binding immunoglobulin protein (Bip), and phosphorylated acetylCoA carboxylase (phsopho-ACC) antibodies were incubated on each slide overnight at 4°C. The slides were then incubated with appropriate secondary antibodies at room temperature for one hour. Staining was visualized using DAB Chromogen before staining with hematoxylin. Images were captured and analyzed using the Motic scanner and ImagePro software (Vista, CA).

### Tumor transcriptomic profile

Transcriptomic analysis from tumor tissues (N=7 mice/group) was performed in the Functional Genomics Core of UNC CH. Briefly, total RNA was isolated from endometrial tumors, sense-strand labeled cDNA was synthesized, and cDNA hybridized to a Mouse Clariom S Human Transcriptome Assay microarray plate (Affymetrix, Santa Clara, CA) and the plate was scanned by a GeneTitan MC Instrument (Applied Biosystems, Waltham, MA). The results were analyzed using Transcriptome Analysis Console Software v 4.0.2 (Thermo Fisher Scientific).

### Tumoral metabolomic profile

Metabolomic profiling was performed on the endometrial tumors from HFD, LFD, HFD-LFD and HFD-IER mice. Metabolites were extracted from 100 mg of tumor tissue (N = 7 mice/group) then analyzed by Metabolon (Research Triangle Park, NC) according to their standard protocol using ultrahigh performance liquid chromatography (Waters Corporation, Milford, MA) coupled with tandem mass spectrometry (UHPLC/MS/MS; Thermo-Finnigan, San Jose, CA), or gas chromatography/MS analysis (Thermo-Finnigan). The changes of tumoral metabolites were analyzed in Omic Insight (Durham, NC).

### Tumoral oxylipinprofile

Oxylipin levels in EC tissue (N=7 mice/group) were analyzed by the Vanderbilt University School of Medicine Eicosanoid Core Laboratory following standard protocols. The extracts from the EC tissues were loaded onto an Oasis MAX micro-elution plate from Waters Corp (Milford, MA). Once eluted, the oxylipins were separated using a Waters Corp Acquity I-Class UPLC and detected using a Waters Corp Xevo TQ-XS triple quadrupole mass spectrometer. To quantify the oxylipins, the ratio of the sample signal to the peak height of an internal isotopically labeled standard was used, and the values were then normalized to the mass of the EC tissue that was used in the analysis.

### Statistical analysis

Results are presented as means ± SD. Statistical significance was evaluated by the unpaired Student’s t-test, one-way ANOVA with Tukey’s multiple comparison test and two-way ANOVA using GraphPad Prism 8 software (La Jolla, CA). P <0.05 were considered statistically significant.

## Results

### IER, LFD and HFD-LFD promote weight loss and inhibits obesity-enhanced endometrial tumor growth in a mouse model of EC

To compare how dietary interventions impact body weight changes and adiposity in the *Lkb1*^*fl/fl*^*p53*^*fl/fl*^ mouse model of EC, mice were randomized into four groups: HFD, LFD, HFD switched to LFD (HFD-LFD), and HFD switched to IER (HFD-IER) (Figure 1A). In line with our prior results(14), after 26 weeks of diet intervention, HFD increased body weight by 33% compared with LFD, resulting in phenotypically obese and lean mice, respectively. Switching from HFD to LFD (HFD-LFD) or IER (HFD-IER) decreased body weight by 27% and 47%, respectively, compared with mice maintained on HFD, achieving weights comparable with or below, respectively, that of lean, LFD-fed mice (Figure 1B). The final weight was lowest in the HFD-IER groups. Since the gonadal fat pad exhibits a strong correlation with body fat(17), we used the ratio of gonadal fat weight to body weight as a measure of adiposity. As expected, the ratio was significantly reduced in the HFD-LFD group and more so in the HFD-IER group, compared with the HFD group, with adiposity comparable with or less than that of the LFD group (Figure 1C).

**Figure 1.**
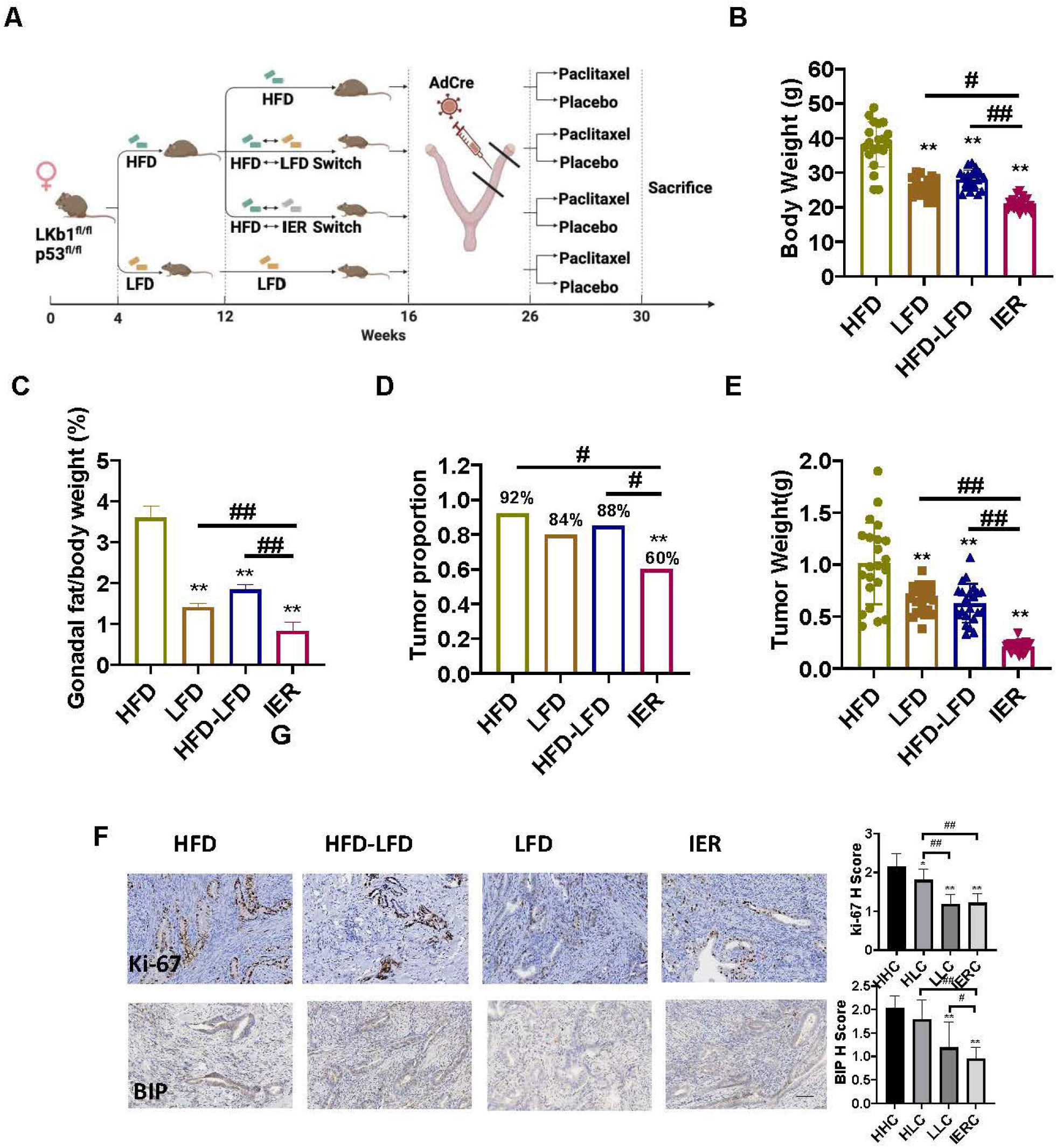
Dietary inventions inhibited obesity-driven EC tumor growth and. Research design diagram. HFD, high fat diet; LFD, low fat diet; HFD-LFD: Changing mouse diet from HFD to LFD; HFD-IER, Switching mouse diet from HFD to IER (A). Weight changes at the end of treatment in each group. HFD-IER significantly reduced the body weight compared to each group (B). Changes in gonadal fat/body weight ratio at the end of treatment in each group. The HFD-IER group showed the most significant reduction compared with the other groups (C). The HFD-IER significantly decreased the incidence of EC in *Lkb1f*^*l/fl*^*p53*^*fl/fl*^ mice compared with mice fed HFD and HFD-LFD, respectively (D). Changes in tumor weight at the end of treatment. The HFD-IER diet was the most effective diet in inhibiting tumor growth compared with each group (E). IHC staining results showed that the HFD-IER diet effectively reduced the expression of Ki67 and Bip compared to the HFD, HFD-LFD and LFD groups (F). Treatment of *Lkb1f*^*l/fl*^*p53*^*fl/fl*^ mice with PTX for 4 weeks did not reduce body weight in each treatment group compared with corresponding untreated mice (G). The HFD-IER mice treated with PTX exhibited the lowest tumor weight compared to HFD, HFD-LFD and LFD mice treated with PTX (H). ^*^ and ^#^<P0.05, ^**^ and ^##^<p0.01. ^*^ and ^**^: compared to HFD group.

To determine the preventive effect of dietary interventions on obesity-related carcinogenesis, we compared the incidence and weight of endometrial tumors in the four groups in the *Lkb1f*^*l/fl*^*p53*^*fl/fl*^ mice14 weeks after AdCre injections. HFD-IER significantly reduced the incidence of EC by 32% and 28% compared with HFD and HFD-LFD, respectively (Figure 1D). Likewise, consistent with our previous studies(14), HFD significantly increased tumor weight, with a 34% increase compared with LFD. Switching from HFD to LFD or IER significantly reversed the obesity-driven increase in tumor weight, with 38% and 79% decreases, respectively, compared with HFD, and both the HFD-LFD and HFD-IER groups were phenotypically formerly obese. The HFD-IER group had the lowest mean tumor weight among all groups (Figure 1E). Tumoral expression of Ki67 and Bip was also significantly decreased in mice fed LFD, HFD-LFD, and HFD-IER diets, when compared to mice fed HFD. The lowest levels of tumoral Ki67 and Bip expression were in the LFD and HFD-IER (phenotypically formerly obese mice via IER. Figure 1F)

### IER, LFD and HFD-LFD enhance the chemotherapeutic effects of PTX in a mouse model of obesity-enhanced endometrial tumor growth

To investigate whether different dietary regimens affect the sensitivity of *Lkb1*^*fl/fl*^*p53*^*fl/fl*^ mice to PTX, each of the diet groups was further randomized 10 weeks after AdCre injections to treatment with PTX (10 mg/kg/week, intraperitoneal) or placebo for 4 weeks. PTX, relative to placebo, did not affect body weight in any diet group (Figure 2A). PTX significantly inhibited tumor growth in HFD, LFD, HFD-LFD and HFD-IER mice, with the inhibition rates in each group ranging from 42.1% to 65.3% (Figure 2B). Compared with other groups, HFD-IER group had the smallest tumors after PTX treatment (0.16g) (HFD 0.33g; HFD-LFD 0.21g; LFD 0.24g). These results suggest that the combination of each dietary regimen with PTX was highly effective in reducing tumor weight in *Lkb1*^*fl/fl*^*p53*^*fl/fl*^ mice, with IER + PTX producing the greatest benefit.

**Figure 2.**
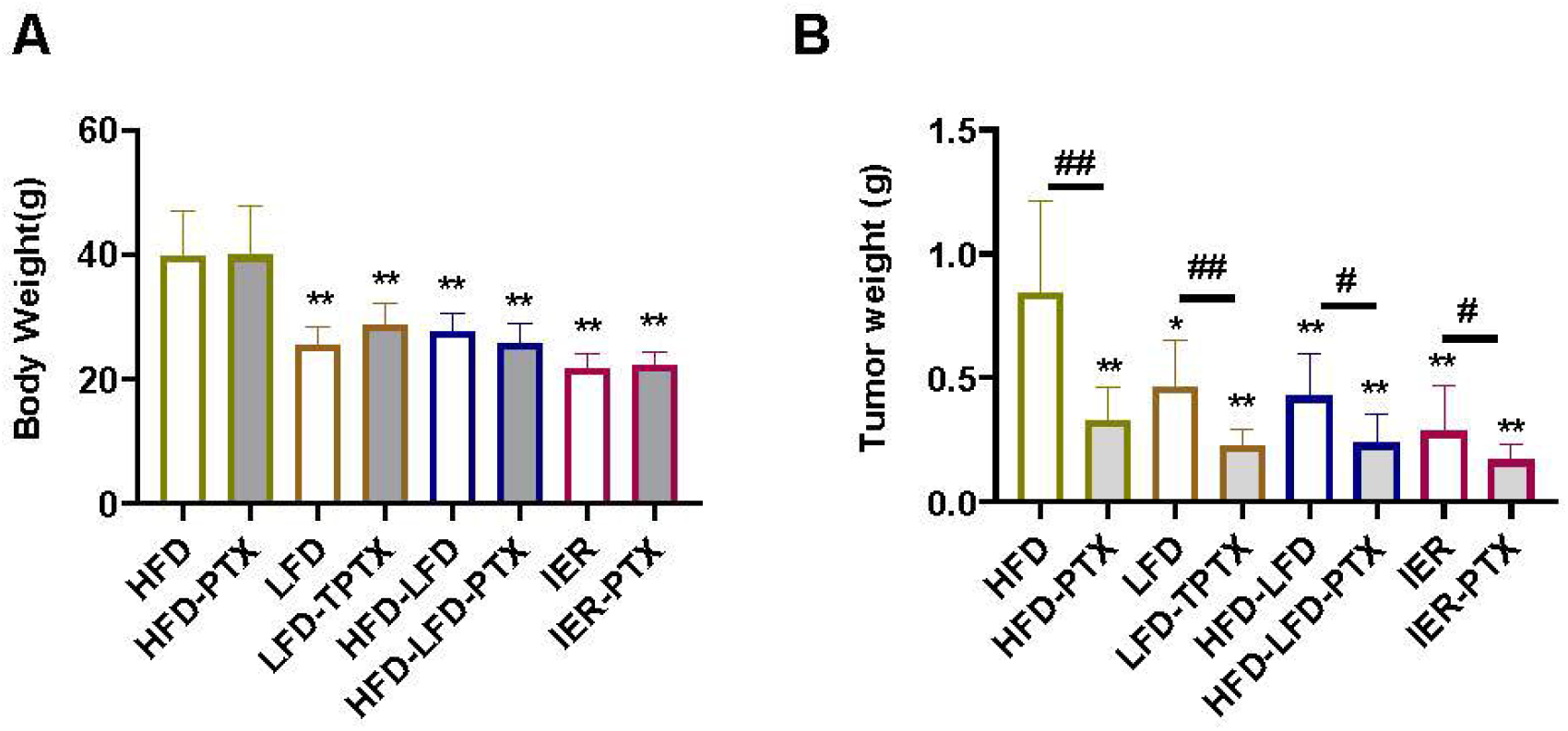
IER increased the sensitivity to PTX. Treatment of *Lkb1f*^*l/fl*^*p53*^*fl/fl*^ mice with PTX or vehicle for 4 weeks did not reduce body weight in each treatment group compared with corresponding untreated mice (A). The HFD-IER mice treated with PTX exhibited the lowest tumor weight compared to HFD, HFD-LFD and LFD mice treated with PTX (B). ^*^ and #<P0.05, ^**^ and ##<p0.01. ^*^ and ^**^: compared to HFD group.

### Dietary regimens differentially affect serum hormones, cytokines and chemokines

Compared with HFD, the LFD, HFD-LFD and HFD-IER groups had significantly reduced serum levels of insulin, leptin and GLP1 (Figure 3A-C). Likewise, the LFD and HFD-IER groups had significantly decreased serum glucagon concentration and increased serum adiponectin concentration, compared with the HFD and HFD-LFD groups (Figure 3D and E). The HFD-IER group had a significantly greater ghrelin concentration than the other groups (Figure 3F). These results indicate that changes in diet enable changes in the systemic hormonal and metabolic environment, and that IER may be a potent way to reverse a HFD-induced protumor environment.

**Figure 3.**
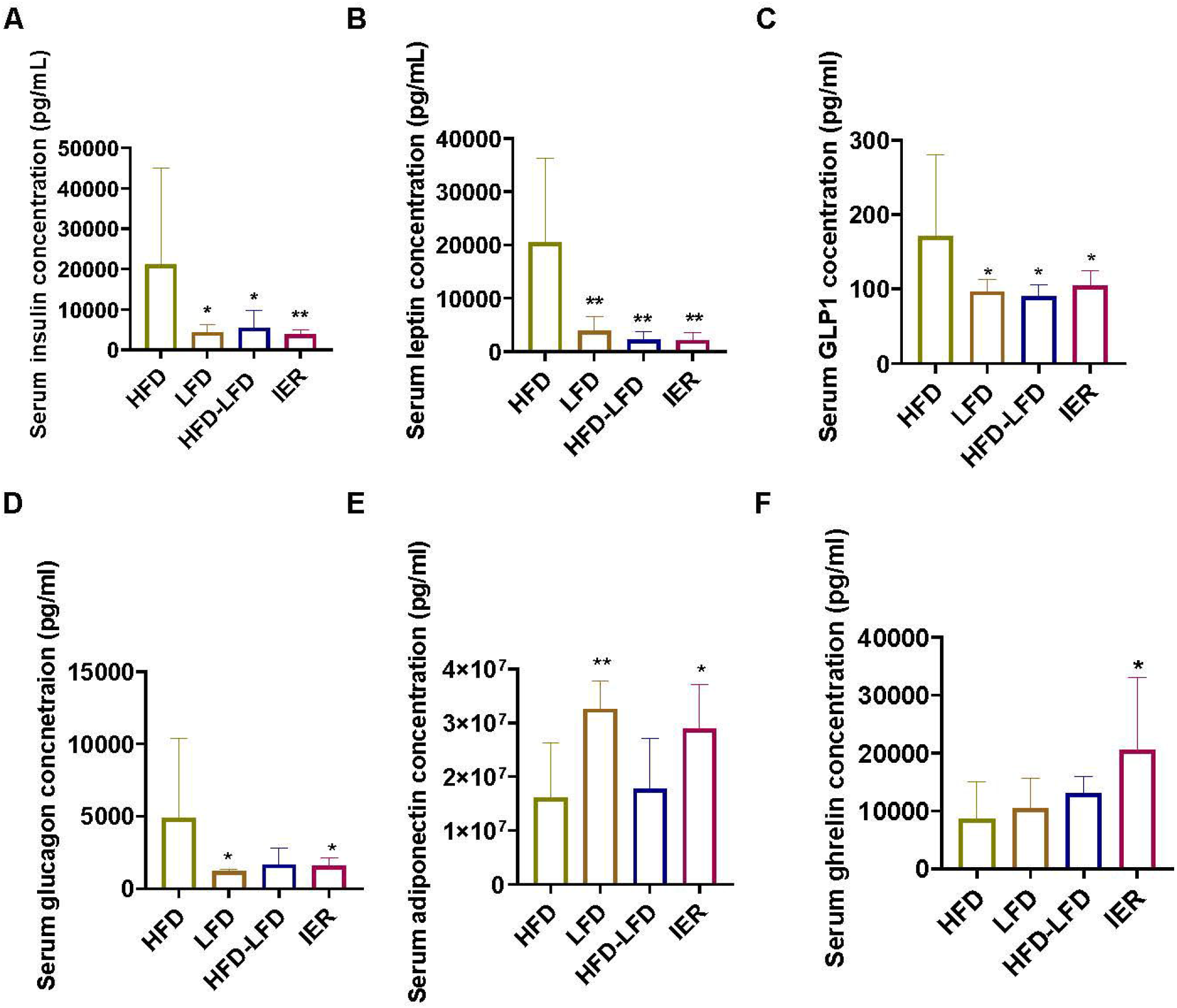
Effects of different dietary regimens on serum hormones in *Lkb1f*^*l/fl*^*p53*^*fl/fl*^ model. Circulating hormones were determined by multiplex ELISA assays. HFD-LFD, LFD and HFD-IER dietary groups had significantly reduced serum concentrations of insulin, leptin, and GLP1, compared to HFD (A-C). LFD and HFD-IER groups had decreased serum concentration of glucagon and increased serum concentration of adiponectin, compared to each group, respectively (D and E). the HFD-IER group had significantly increased serum concentrations of ghrelin compared to the HFD, HFD-LFD and LFD groups (F). ^*^<P0.05, ^**^<p0.01.

Since weight loss may improve the inflammatory environment in EC(3, 18), we next examined changes in a panel of inflammatory cytokines and chemokines in the serum of *Lkb1*^*fl/fl*^*p53*^*fl/fl*^ mice on the different dietary regimens. A heatmap displays the relative serum concentrations of pro-inflammatory cytokines based on diet regimen. The pattern of pro-inflammatory cytokine concentrations in the HFD-IER group closely resembled that of the LFD (Figure 4A). IL-2, IL-4, IL6, and IL-16 levels were significantly decreased in the LFD and HFD-IER groups compared with the HFD group, and there was no difference between the HFD and the HFD-LFD groups (Figure 4B-E), suggesting that an inflammatory environment still exists in HFD-LFD mice despite being phenotypically formerly obese. Similarly, MIP-3b, RANTEs (CCL5), SCYB 16 (CXCL 16), and TNFα levels followed the same pattern by diet group as did the interleukin family, including significant decreases in LFD and HFD-IER mice compared to HFD mice (Figure 4F-I). Additionally, only HFD-IER mice had lower MCP 6 concentrations compared with HFD and HFD-LFD mice (Figure 4J). These results showed that IER mitigated obesity-related perturbations in serum cytokines and chemokines to such an extent that levels become comparable to those in lean, never obese mice.

**Figure 4.**
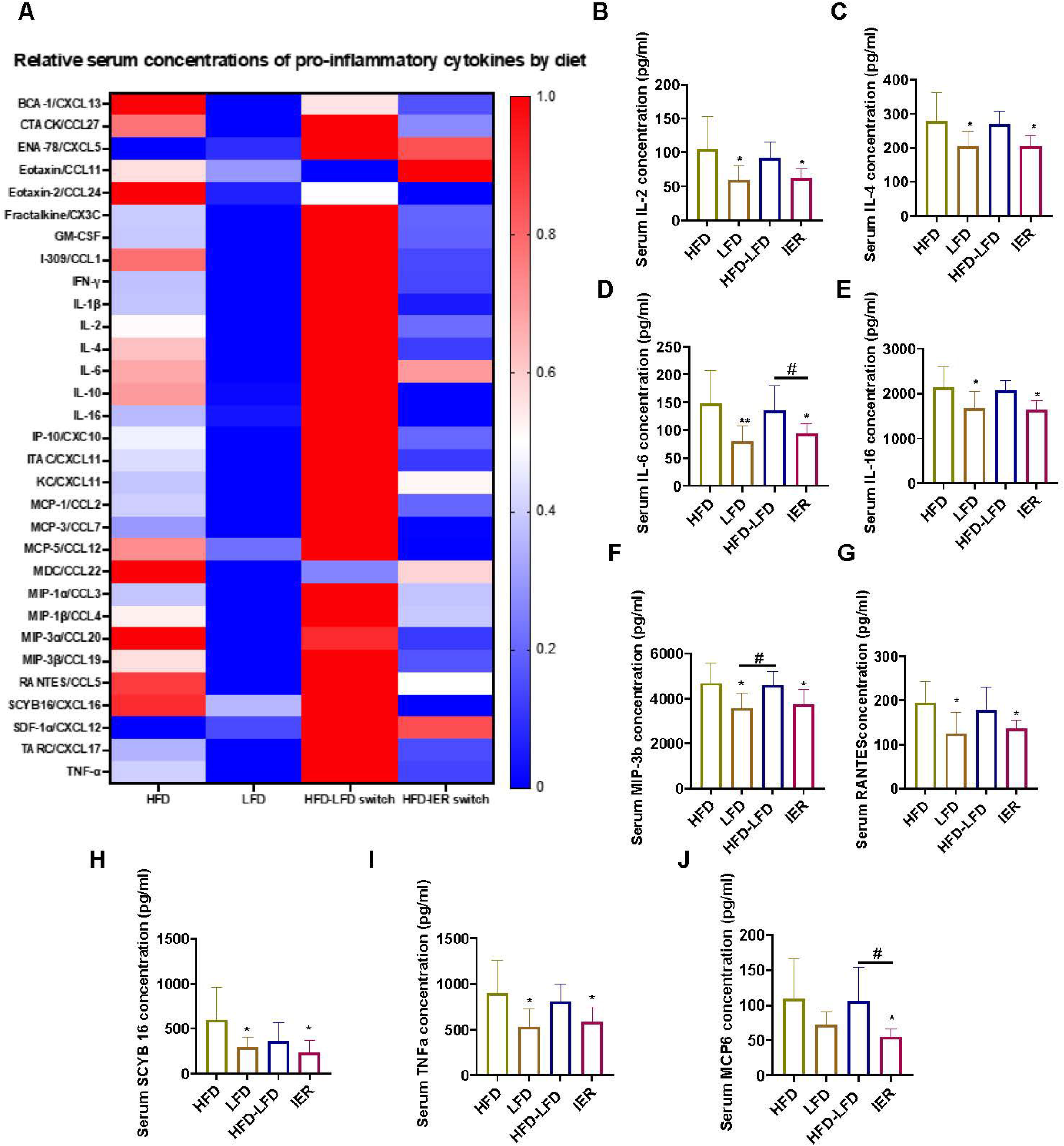
Different dietary regimens affect changes in cytokines and chemokines. Circulating cytokines and chemokines were determined by multiplex ELISA assays. The heat map showed that dietary regimens differentially altered the changes pro-inflammatory cytokines (A). The LFD and HFD-IER mice had significantly decreased serum IL-2, IL-4, IL6, and IL-16 levels compared with the HFD mice (B-E). Similarly, the LDF and HFD-IER mice had reduced serum concentrations of MIP-3b, RANTEs (CCL5), SCYB 16 (CXCL 16), and TNFα compared with the HFD mice (F-I). The HFD-IER mice had the lowest serum concentrations of MCP 6, compared with HFD and HFD-LFD mice (J). ^*^ and ^#^<P0.05, ^**^<p0.01. ^*^ and ^**^: compared to HFD group.

### Dietary regimens differentially affect tumoral gene expression profiles

To investigate the changes in global gene expression profiles of EC tissues with different dietary regimens, we identified differentially expressed genes in HFD, LFD, HFD-LFD and HFD-IER groups using transcriptomic profiling of tumors via Affymetrix microarray, followed by gene set enrichment analysis (GSEA). There were significantly different gene expression profiles in each group. The HFD group had enriched tumoral expression in estrogen response, inflammatory response, adipogenesis, and cholesterol metabolism, while the LFD group had improved inflammatory responses and enriched expression of G2M checkpoint, E2F and MYC targets, DNA repair, oxidative phosphorylation, unfolded protein response, and glycolysis in comparison of these two groups(Figure 5A). In comparison between the HFD and HFD-LFD groups, the HFD-LFD regimen significantly revised HFD-induced activities of tumoral estrogen responses, cholesterol homeostasis, and K-ras and p53 signaling pathways (Figure 5B). Likewise, compared with the HFD-IER group, the HFD group had higher tumoral inflammatory and immune responses and activated E2F pathway, while the HFD-IER regimen effectively revised these inflammatory and immune responses and increased the functions of adipogenesis, apical Junction, hedgehog signaling and hypoxia pathways (Figure 5C). Additionally, we have compared HFD-LFD vs LFD, HFD-IER vs

**Figure 5.**
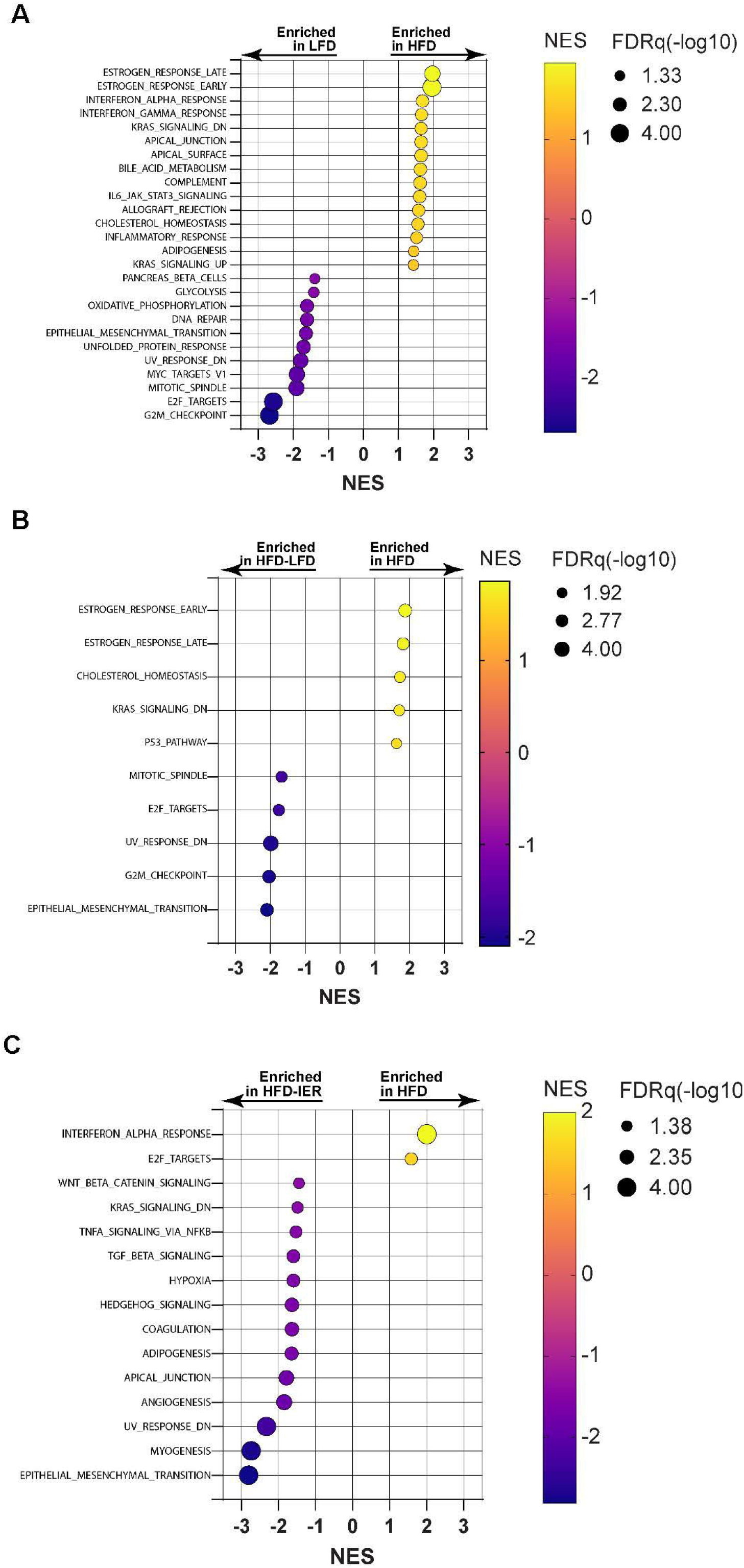
Transcriptomic analysis of tumor tissues at the end of dietary intervention. Significant signature gene sets were determined by pairwise comparison of transcriptomics of endometrial tumor tissue from *LKB1*^*fl/fl*^ *p53*^*fl/fl*^ mice with HFD by GSEA. LFD vs HFD (A), HFD-LFD vs HFD (B), and HFD-IER vs HFD (C). Biological processes were ranked as enrichment scores for each gene set. The color of the bubbles represents the enrichment fraction. The size of the bubble represents -log10 (P value). n=6/group.

LFD and HFD-IER vs HFD-LFD groups and found that these diet changes also significantly contributed to functional pathway changes in inflammation, immune, estrogen response, glycolysis, cholesterol metabolism, and DNA repair (Supplemental figure 1).

### Dietary regimens differentially affect tumoral metabolic and oxylipin profiles

To determine changes in biochemical and metabolic pathways in mice on different dietary regimens, we analyzed changes in metabolites in EC tissues from all four groups by Metabolon assay. By metabolic profiling, 816 biochemicals were identified and analyzed in all EC tissues. Of these 816 biochemicals, 243 differed between the HFD and LFD groups, with 212 upregulated and 31 downregulated. Many of the metabolites affected were amino acids, lipids, nucleotides, and peptides, including triacylglycerols (TAGs), diacylglycerol (DAG), fatty acids and cholesterol ester lipid classes, which were all increased in the HFD group compared with the LFD group (Supplemental Figure 2). Compared with the HFD group, the LFD and HFD-IER groups had significantly reduced tumoral metabolism of amino acids, nucleotides, nicotinate, ascorbate, and aldarate(Figure 6A). Similarly, significant reductions in cholesterol esters, diacylglycerol and hexosylceramide were found in the tumors of HFD-LFD, LFD, and HFD-IER mice, with tumors from HFD-IER mice showing the most dramatic reduction (Figure 6B). Tumoral oxidative stress markers (ophthalate and 4-hydroxynonenal glutathione) were significantly reduced in LFD and HFD-IER mice (Supplemental Figure 3). In HFD-IER mice, fatty acids (including palmitic acid, lauric acid, myristic acid, and docosahexaenoic acid) and β-oxidation were increased in tumor tissues compared to other groups, indicating that the metabolism of fatty acids is involved in the inhibition of tumor growth in phenotypically formerly obese (HFD-IER) mice (supplemental Figure 4). Overall, these results indicate that switching an obese mouse from HFD to either LFD or IER modifies the obesity-induced environment to achieve an environment similar to that of never-obese mice (LFD), and IER appears to be more effective than LFD in this regards.

**Figure 6.**
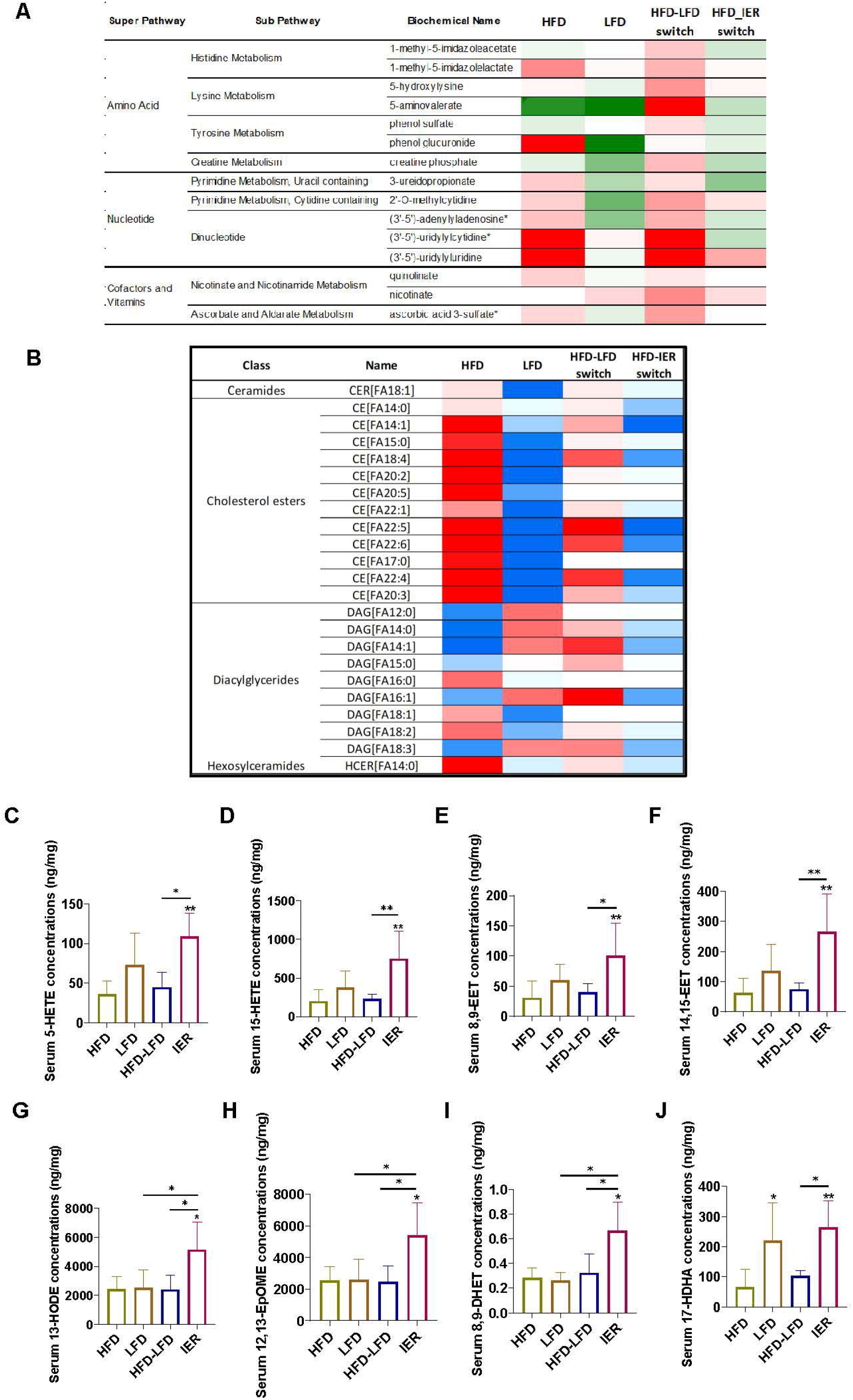
Effects of different dietary regimens on tumoral metabolic and oxylipin profiles. Metabolomic analysis results showed that HFD-IER dietary intervention significantly reduced the metabolism of amino acids, nucleotide amino acids, nucleotides, nicotinate, ascorbate, and aldarate in endometrial tumor tissues, compared to HFD, HFD-LFD and LFD (A). HFD-LFD, LFD, and HFD-IER regimens decreased the tumoral metabolism of cholesterol esters, diacylglycerol and hexosylceramide, with HFD-IER exerting the greatest effect compared to other groups (B). Eight oxylipin molecules from different PUFAs were analyzed. Compared to tumors from HFD and HFD-LFD mice, tumors from HFD-IER mice had significantly increased the concentrations of 6-HETE, 15-HETE, 8,9-EET and 15,15-EET (C-F). Compared with tumors from HFD, HFD-LFD and LFD mice, tumors for HFD-IER mice had a significant increase in 13-HODE, 12, 13 EpOME, and 8,9 DHET (G-I). LFD and HFD-IER mice had a significantly increased the concentration of 17-HDHA in tumor tissues compared to HFD mice (J). N=7/group, ^*^<P0.05, ^**^<p0.01.

Since endometrial tumors in HD-IER mice have active fatty acid metabolism and oxylipins from the enzymatic or non-enzymatic oxidation of polyunsaturated fatty acids (PUFAs), we compared changes in tumor tissue oxylipins among the diet groups. Eight oxylipin molecules from different PUFAs were significantly increased in tumors from the HFD-IER group compared with the HFD-LFD and the HFD groups (Figure 6 C-J). Compared with the HFD, HFD-LFD and LFD groups, the HFD-IER regimen had a significant increase in tumoral 13-HODE, 12, 13 EpOME, and 8,9 DHET (Figure 6G-I). Tumoral 17-HDHA, an autoxidation product of docosahexaenoic acid (DHA), was significantly increased in the LFD and HFD-IER groups compared with HFD group (Figure 6J). These results indicate that oxylipins are involved in weight loss and inhibition of tumor growth in HFD-IER mice.

## Discussion

This study presents a comprehensive analysis comparing the impact of different dietary regimens (i.e., HFD, LFD, HFD-LFD and HFD-IER), with and without PTX, on body weight, tumor growth, cytokines, hormones, gene expression and metabolic parameters in a transgenic mouse model of EC under obese and lean conditions. Overall, we conclude that weight-reducing dietary interventions in obese mice, specifically, switching from HFD to either LFD or (even more so) IER, decreases body weight and adiposity; reverses obesity-associated perturbations in serum hormones, cytokines and chemokines; reverses obesity-associated, protumorigenic transcriptional and metabolic profiles in the tumor environment; and particularly when in combination with PTX, effectively reduces tumor incidence and progression in a mouse model of EC.

Women with weight loss, especially intentional weight loss in postmenopausal obese women, have a significantly lower endometrial cancer risk(5). Dietary intervention is the first line of treatment for obese women, which involves restricting energy intake. IER regimens have positive impacts on body weight, metabolic profiles, inflammation, oxidative stress, and insulin sensitivity in both animal models and humans(19). However, in some preclinical models, research results on the role of IER in tumor prevention and treatment are inconsistent. The length of IER treatment, changes in body weight and insulin, different IER regimens, and unknown genetic factors might influence the outcomes of IER in animal models of cancer(8, 20). We compared the effects of four dietary interventions on tumorigenesis and tumor growth in *Lkb1*^*fl/fl*^*p53*^*fl/fl*^ mice in the current study and found that HFD-induced obesity significantly increased endometrial tumor growth, while maintaining a lean phenotype (via continuous LFD) or reversing obesity (via LFD or IER) all effectively reduced tumor weights relative to the HFD. The anticancer effects of LFD and IER were accompanied by reduced serum insulin, leptin, GLP1 and glucagon levels, and increased ghrelin, compared with HFD mice. Importantly, IER was generally more effective in reducing body weight, obesity-rlated hormone levels, and development and progression tumor of EC than LFD,, suggesting IER may be a more effective weight loss and cancer prevention strategy than LFD for EC in patients with overweight/obesity.

In some animal models and clinical trials, fasting combined with chemotherapy significantly protected normal cells from the drug’s harmful side effects, reduced their toxicity, and improved the efficiency of chemotherapy(21, 22). Fasting 48 hours before chemotherapy effectively inhibited tumor growth compared to chemotherapy alone in fibrosarcoma mouse models(13). However, in a colorectal cancer mouse model, fasting combined with chemotherapy did not improve tumor response, although the fasting mice experienced fewer side effects(23). Our results demonstrate that PTX effectively inhibited tumor growth in *Lkb1*^*fl/fl*^*p53*^*fl/fl*^ mice under different dietary inventions without obvious side effects, and with particular benefit when used in combination with IER.

The increased adiposity accompanying obesity results in adipose-derived metabolic changes, including the overproduction of pro-inflammatory mediators and changes in adipokine secretion (including increases in leptin) associated with insulin resistance, immunosuppression and increased development and progression of EC(24, 25). IER reduced markers of inflammatory responses and oxidative stress in DSS-induced colon tissues, and decreased serum levels of oxidative and inflammatory factors such as lipopolysaccharide, malondialdehyde, TNF-α and IL-6 in C57BL/6 mice(26). Twelve weeks of IER may be beneficial in reducing obesity-related inflammation and insulin resistance, regardless of weight loss, compared with a continuous energy restriction in a randomized controlled clinical trial (27). A recent meta-analysis demonstrated IER is more effective at reducing serum C-reactive protein levels than continuous energy-restricted diets, with more pronounced effects in overweight and obese individuals(28). We analyzed 30 serum cytokines and chemokines in HFD, LFD, HFD-LFD and HFD-IER groups and found that HFD-IER significantly reduced IL-2, IL4, IL-6, IL16, MCP6, MIP-3b, RANTES, SCYB 16 and INF-α levels to those in LFD mice, indicating that IER effectively reduces obesity-driven inflammatory responses in *Lkb1*^*fl/fl*^*p53*^*fl/fl*^ mice.

CR or IER can alter the expression of genes and metabolites and produce beneficial effects by increasing insulin sensitivity and decreasing serum levels of triglycerides and low-density lipoproteins in animal models and humans(19, 29-32). The IER of 23 overweight premenopausal women at high risk for breast cancer during one menstrual cycle showed that serum and urine metabolites fluctuated significantly, and the gene expression changes caused by IER were more subtle and variable than those caused by sustained ER(33). We found that different dietary regimens manifested different gene expression and metabolite profiles in EC tumor tissues, suggesting that the anticancer effects of these dietary interventions depend on distinct cell signaling pathways in the *Lkb1*^*fl/fl*^*p53*^*fl/fl*^ mice. This conclusion is also supported by our analysis of 816 tumoral metabolites that identified diet regimen-specific changes in metabolites.

Compared with tumors from mice that were obese (HFD) or that were phenotypically formerly obese via LFD (HFD-LFD), tumors from mice that were phenotypically formerly obese via IER (HFD-IER) appears to be more like those from lean, never obese mice (LFD) in terms of altering obesity-induced metabolic and transcriptional changes, including promoting an inflammatory environment, reducing metabolism of amino acids, nucleotides, and cholesterol, increasing β-oxidation, and reducing oxidative stress in endometrial tumors. It remains unclear if the tumor environment of the HFD-LFD mice would undergo metabolic and/or transcriptional reprogramming to that observed for the HFD-IER and LFD mice after a longer period of sustained weight loss.

Oxylipins are bioactive lipid metabolites derived from PUFAs via the cyclooxygenase, lipoxygenase, and cytochrome P450 pathways and are involved in a wide range of functions including inflammation, immunity, and tumor growth (34, 35). Dietary interventions significantly influence the levels of oxylipins in multiple mouse models(36, 37). For the first time, we report diet-dependent changes in oxylipins in the EC tissues of *Lkb1*^*fl/fl*^*p53*^*fl/fl*^ mice. Specifically, HFD-IER effectively increased the levels of linoleic acid (LA) derived 12-HODE and 12,13-EpOME, eicosapentaenoic acid (EPA) derived 8,9 DHET, and docosahexaenoic acid (DHA) derived 17-HDHA, as compared with HFD, LFD and HFD-LFD groups; and increased the products of arachidonic acid derived 5-HETE, 15-HETE, 8,9-EET and 14,15-EET, as compared with HFD and HFD-LFD mice. Given the anti-inflammatory effects of arachidonic acid and our recent finding that LA, DHA, and EPA significantly inhibited cell proliferation and tumor growth in EC cell lines and the *Lkb1*^*fl/fl*^*p53*^*fl/fl*^ mouse model of EC(38), we speculate that IER may exert both beneficial anti-inflammatory and antitumorigenic activities through elevated oxylipins.

## Conclusions

IER and (less so) LFD, alone and in combination with paclitaxel, promote weight loss, reverse the procancer effects of HFD-induced obesity, and mitigate tumor development and progression in a mouse model of EC. These findings lay the foundation for further investigations of IER as an EC prevention and treatment strategies, potentially in combination with PTX, in women with overweight/obesity.

## Data availability

All data generated or analyzed during this study are included in this article. The datasets used and/or analyzed during the current study are available from the corresponding authors upon reasonable request.

## Author Contributions

ZZ, ZF, WCB, MAN, BD, XS, HS and WS performed experiments. ZZ, MC analyzed microarray. GM detected oxylipins. DL analyzed metabolomic data. SH provided resources and revised manuscript. JW, WK, and CZ analyzed and interpreted the data. ZZ, CZ and VB prepared the manuscript. CZ and VB designed the experiments, revised the manuscript, and provided financial support. All authors have read and approved the final manuscript.

## Ethics approval

All animals’ experimental procedures were approved by Institutional Animal Care and Use Committee of University of North Carolina at Chapel Hill (protocol #21-229).

