## Supplementary figures and images for "Intermittent energy restriction inhibits tumor growth and enhances paclitaxel response in a transgenic mouse model of endometrial cancer"

### Supplemental Figure 1

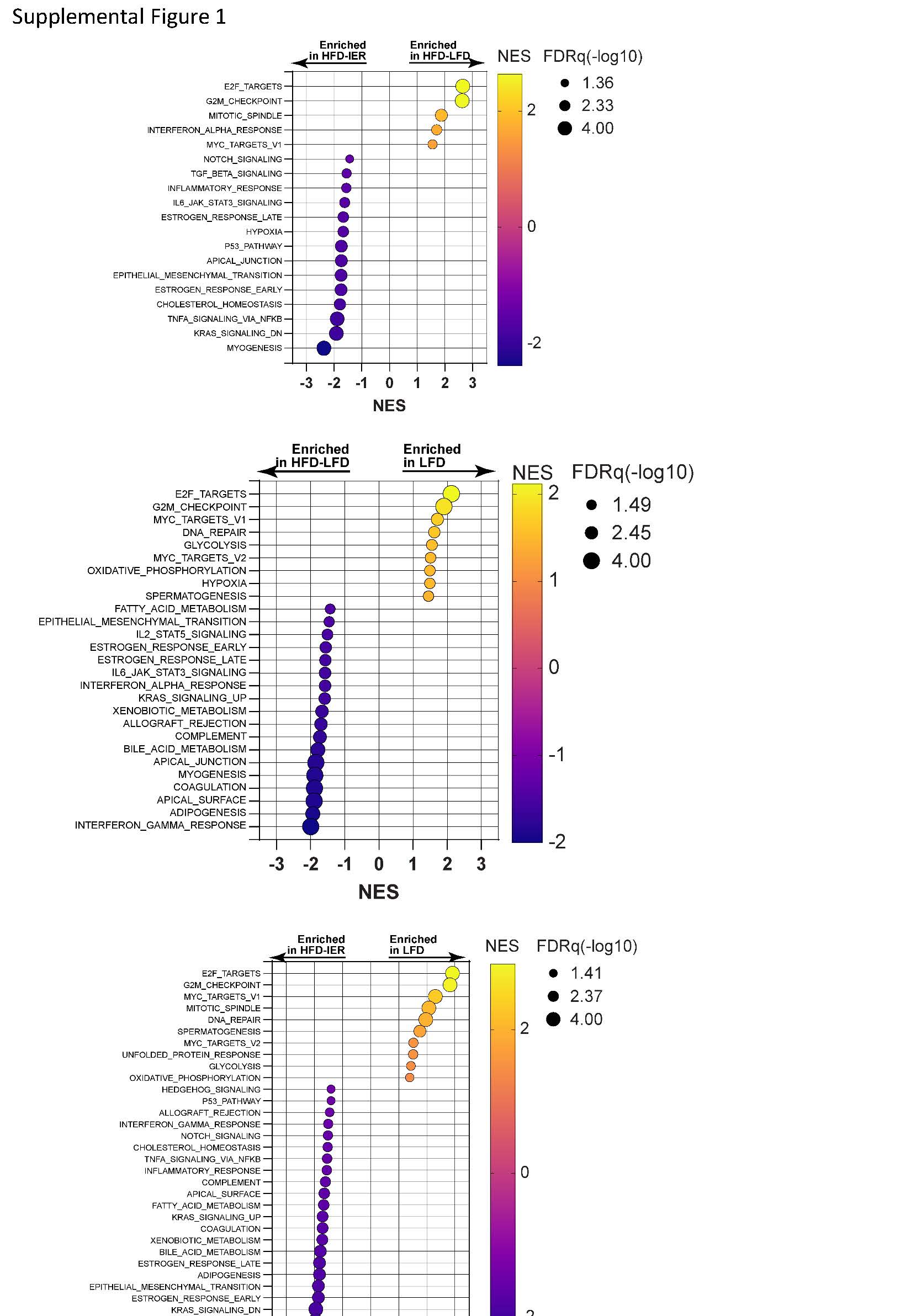

### Supplemental Figure 2

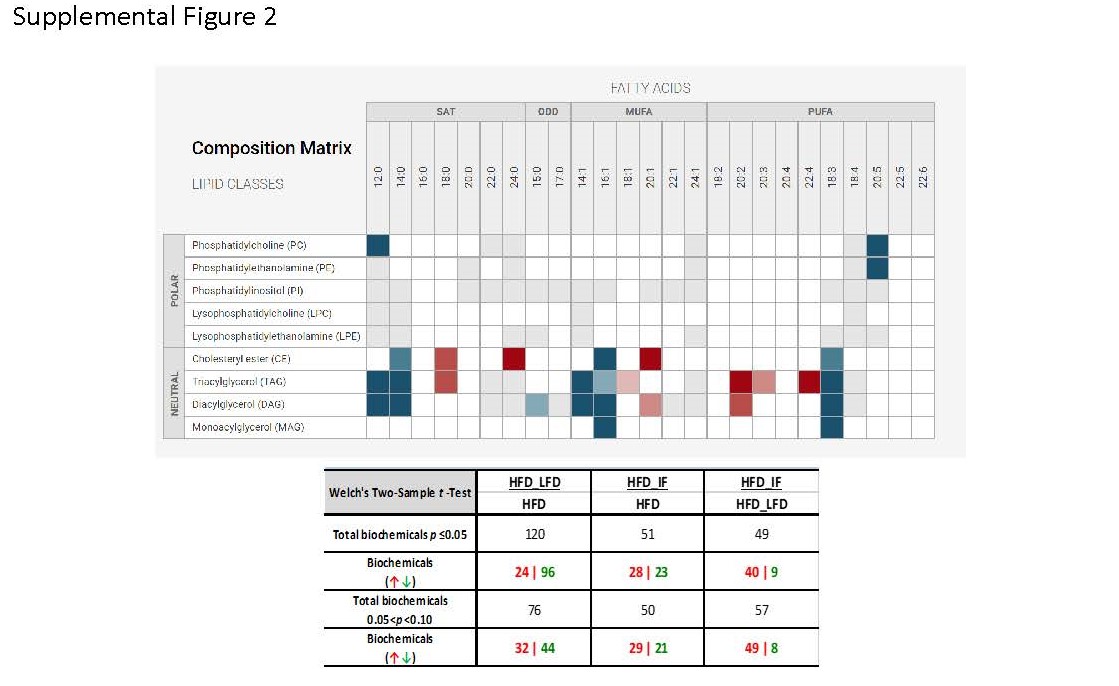

### Supplemental Figure 3

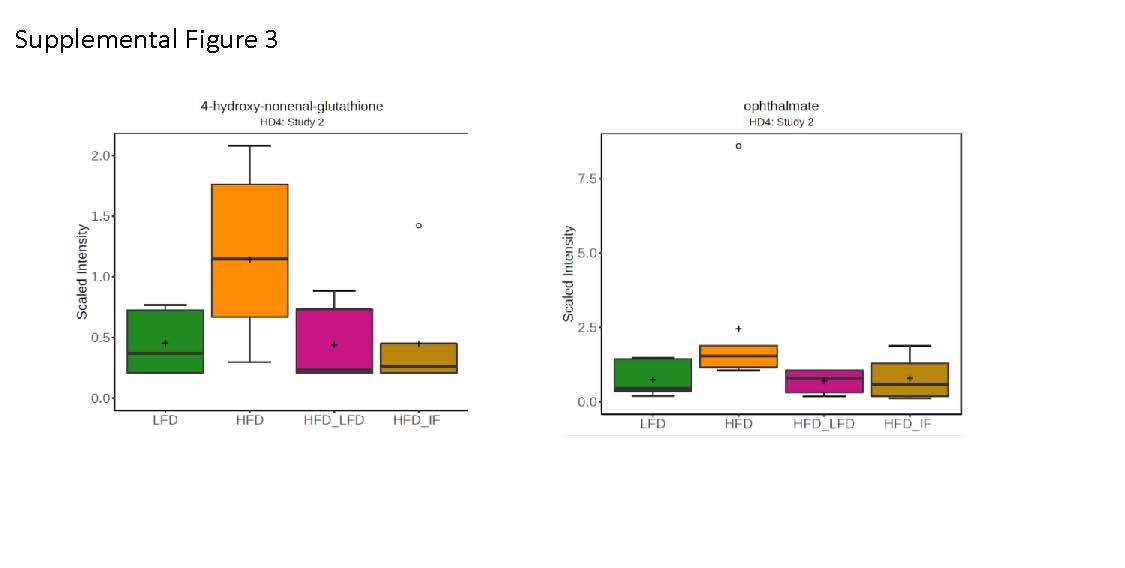

### Supplemental Figure 4

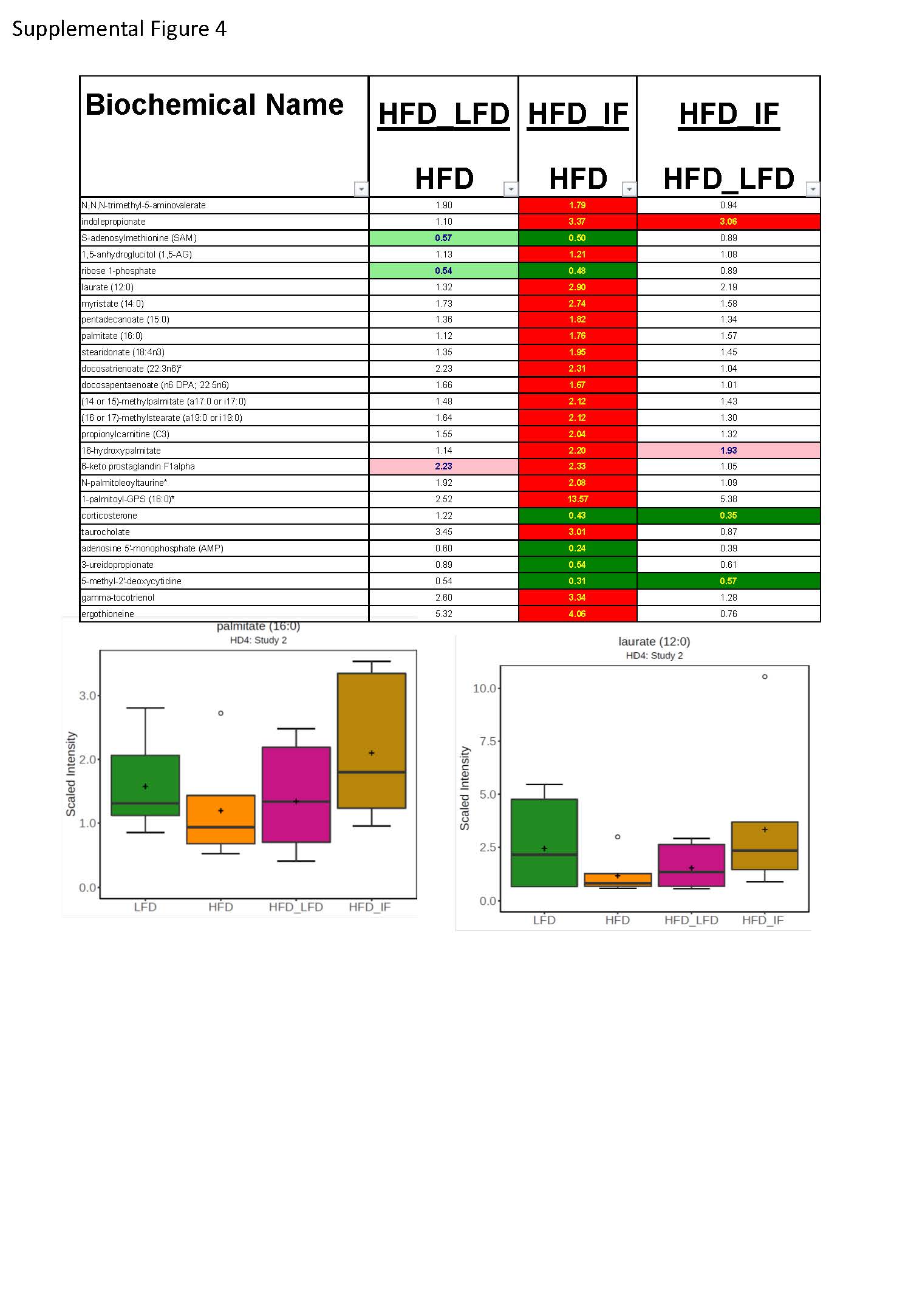
